# Mitochondrial complex I controls blood brain barrier permeability

**DOI:** 10.1101/2022.10.13.512023

**Authors:** Gavin M. Davis, Estelle Juere, Jerrard M. Hayes, Gavin P. Davey

**Affiliations:** School of Biochemistry and Immunology, Trinity College Dublin, Dublin 2, Ireland; BioCyto Ltd, Carragaline, Co Cork, Ireland

**Keywords:** Blood brain barrier, mitochondria, neurodegeneration, human induced pluripotent stem cells, metabolism, complex I, bioenergetics

## Abstract

Mitochondrial electron transport chain (ETC) complexes are key mediators of energy metabolism in astrocytes and neurons, with subsequent effects on memory, behaviour and neurodegeneration. Mitochondrial dysfunction and increased blood brain barrier (BBB) permeability are known pathologies in Parkinson’s and Alzheimer’s diseases. However, knowledge of how ETC activities regulate metabolic flux and influence permeability in the BBB is lacking. Using metabolic flux control analysis we show that complex I is a critical control point for oxidative flux and permeability in brain microvascular endothelial cells derived from human induced pluripotent stem cells. Inhibition of complex I activity immediately reduced the transendothelial electrical resistance (TEER) by 60%, leading to an increase in protein transport across the BBB. These events were accompanied by a transient reduction in ATP that was recovered, along with TEER values, over an extended time period. Furthermore, while inhibition of downstream complexes III or IV decreased oxygen respiration rates, no effects on BBB permeability were identified, due to compensatory glycolytic flux and maintenance of ATP synthesis. These data show that mitochondrial complex I is critical for maintaining energy production in endothelial cells and transiently controls BBB permeability, which may contribute to brain disorders where complex I dysfunction is a hallmark.

## Introduction

The blood brain barrier (BBB) is a specialized cellular barrier composed of unique brain microvascular endothelial cells (BMECs). The BBB is critical in the transport of metabolites and solutes into the brain, while preventing unregulated transport of blood born toxins, breakdown products, proteins and immune cells^1^. BMECs create the BBB by expression of specialized tight junction proteins and junctional adhesion molecules which allow tight intercellular contacts, known as tight junctions^2,3^. Increased blood brain barrier (BBB) permeability is a well characterised pathology in many neurodegenerative disorders^4^.

Complex I (CI-NADH:Ubiquinone oxidoreductase) is a 45 subunit protein complex located in the mitochondrial inner membrane (MIM). Electrons transferred from NADH to CI are shuttled through the electron transport chain (ETC), contributing to a proton-electrochemical gradient that is harnessed by ATP synthase for the production of ATP. As complex I is a key regulator of energy metabolism in the brain, affecting both memory and behaviour^5–7^, dysfunction of the complex is considered an important factor in neurodegenerative disorders^1^.

In line with this, the activity of CI is decreased by 30-40% in the substantia nigra of post mortem brain samples from Parkinson’s disease (PD) patients^8^, with decreased activity detected in mitochondria from other brain regions such as the frontal cortex^9^. Mutations in the ND5 subunit of CI have been detected in the PD brain indicating that dysfunctional CI may be present in mitochondria throughout the brain and peripheral tissues^10^. CI and complex IV (CIV) dysfunction have also been reported in Alzheimer’s disease (AD), with decreased CI activity leading to a deficit of mitochondrial respiration, reduction of ATP synthesis and increased pathological reactive oxygen species (ROS)^11^.

There is a growing body of evidence highlighting BBB dysfunction in pathologies such as PD and AD with findings such as altered tight junction protein complexes, increased permeability and altered function of efflux transporters such as P-glycoprotein ^10,11^. Here we used BMECs derived from iPS (IMR90)-4 cells ^12,13^ to evaluate the contribution of electron transport chain dysfunction to BBB dysfunction. Metabolic Control Analysis (MCA) was employed to determine the level of control complex I has over mitochondrial respiration, indicating a flux control co-efficient of 0.25 ± 0.07 in BMECs and an inhibition threshold of approx. 52%. This in depth metabolic analysis highlights that complex I, rather than complex III or IV, is a critical control point in BBB permeability and transport of proteins across *in vitro* BBB models. Our study highlights that mitochondrial complex I dysfunction and subsequent metabolic defects present in neurological disorders such as PD and AD could have a role in increasing BBB permeability in the brain.

## Methods and Methods

### Antibodies and Reagents

ZO-1, Claudin-5, OPA1 antibodies were purchased from Abcam, Occludin from Sigma Aldrich, β-Actin from Cell Signalling Technologies, and Mff from Proteintech. Accutase, Collagen IV, bFGF, ROCK inhibitor were purchased from Sigma Aldrich. Human iPS(IMR90)-4 pluripotent stem cells were purchased from WiCell. Growth factor reduced Matrigel was purchased from Corning Ltd. KnockOut serum replacement (KOSR), human endothelial serum free media (hESFM), B27 supplement and Versene were obtained from Biosciences Ltd. mTeSR1 medium was purchased from Stemcell Technologies.

### *In vitro* generation of brain microvascular endothelial cells from human induced pluripotent stem (iPS) cells

Human induced pluripotent stem cells were cultured and maintained using the Stem Cell Technologies Matrigel mTeSR system, according to WiCell’s feeder independent protocols. BMECs were generated using the method described by Lippman *et al*. (2012)^12^ which uses the human induced pluripotent stem cell line iPS(IMR90)-4 derived from IMR90 cells. iPS(IMR90)-4 cells (WiCell) were fully differentiated into BMECs in 6 well plates over 11 days. On day minus 3 cells were seeded at approx. 0.1 × 10^6^ cells per well on matrigel coated plates in mTeSR1 media and expanded for 3 days. On Day 0 cells from one well were detached using versene according to the manufacturer’s instructions, counted and plated at a differentiation density between 0.3 × 10^6^ per well. mTeSR1 media was replaced with unconditioned media (UM/DMEM F12 with HEPES, KOSR (20%), non-essential amino acids (1%), β-mercaptoethanol (100 μM). Cells were maintained in UM for 6 days with media change every day. On day 6 media was changed to endothelial cell medium (hESFM, Biosciences Ltd) supplemented with 20 ng/mL b-FGF, 10 μM retinoic acid and 0.5% B27 (EC + RA media). Cells were maintained in EC + RA medium for 2 days without any media change. At this stage cells were fully differentiated into BMECs and sub-cultured onto 6-well plates for immunoblotting or transwell filters (3 μm, Millipore) for TEER analysis following singularisation of cells using accutase for 45 minutes and counting. TEER values were taken using a Millicell ERS-2 Voltohmmeter. Transwell filters were previously coated overnight at 37°C using a solution of 4 parts collagen IV (1 mg/mL, Sigma), 1 part fibronectin (1 mg/mL, Sigma) and 5 parts MiliQ water (4:1:5 solution). The coating solution was removed and the transwells air-dryed for 20 minutes in a laminar flow hood and rehydrated with EC media (500 μl). Singularised cells were counted and approx. 1.5 × 10^6^ cells were added to each transwell. Sub-culturing onto plates depended on the type of experiment. For Seahorse XF analysis, 0.1 × 10^6^ cells were plated onto each well of a 96 well Seahorse tissue culture plate. For membrane potential experiments 0.5 × 10^6^ to 1 × 10^6^ cells were plated onto 4 quadrant confocal dishes (Ibidi). A 1 in 5 dilution of 4:1:5 coating solution was used for sub-culturing cells onto plates. 24 h after cells were subcultured onto plates or transwell filters EC + RA media was replaced with EC medium containing only B27 supplement (EC + B27 media). Following another 24 h TEER values reached their maximum and experiments were carried out. Human induced pluripotent stem cells were cultured and maintained using the Stem Cell Technologies Matrigel mTeSR system, according to WiCell’s feeder independent protocols. BMECs were generated using the method described by Lippman *et al*. (2012) which uses the human induced pluripotent stem cell line iPS(IMR90)-4 derived from IMR90 cells.

### Measuring Trans-Endothelial Electrical Resistance (TEER)

TEER measurements were taken using a Millicell ERS-2 Voltohmmeter with an adjustable chopstick electrode. TEER measurements were taken in 3 positions of the transwell chambers by submerging the long arm of the chopstick electrode in the basolateral chamber and the short arm in the apical chamber, according to the manufacturer’s instructions. TEER values consistently averaged between 1500-2500 ohms/cm^2^.

### Confocal Microscopy

Cells (for fixed and live imaging) were imaged using an Olympus FV1000 Point Scanning Confocal Microscope, FV10-ASW Olympus Fluoview Ver. 2 software and a 63 x oil immersion objective and a Leica TCS SP8 STED microscope with associated software and 63 x oil objective. The imaging chamber was heated to 37°C and humidified in an environment of 5% CO_2_ for live cell imaging. Mean fluorescence intensity (MFI) of tetramethylrhodamine, methyl ester (TMRM) stained mitochondria was measured using Imaris imaging software.

### Seahorse XF Analysis

For metabolic analysis of differentiated BMECs (day 8) cells were sub-cultured onto Seahorse cell culture plates and allowed to proliferate for 2 days prior to metabolic analysis. Cells were incubated in a minus CO_2_ incubator for 1 h before performing extracellular flux analysis. Oligomycin (1 μM), FCCP (1 μM) and rotenone (1 μM) / antimycin A (2 μM) were used to calculate proton leak, maximal respiration and non-mitochondrial respiration, respectively.

### ETC complex I assay

The complex I specific activity assay was performed as described in Telford et al., (2009), by following the oxidation of NADH at 340 nm with decylubiquinone as the electron acceptor, using a multichannel Shimadzi UV-VIS spectrophotometer UV-2600. A SpectraMax M3 96-well plate reader was used to determine protein concentration and cellular viability using an alamar blue assay (Invitrogen).

### Metabolic Control Analysis (MCA)

MCA of BMECs was performed by titrating the complex I inhibitor rotenone (0 – 1 μM) during the complex I assay and Seahorse analysis. Seahorse XF analysis was used to measure oxygen consumption rate (OCR) in response to increasing concentrations of rotenone. The flux control co-efficient for complex I was calculated using the following equation, at low rotenone concentrations:

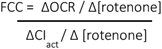

ΔOCR/Δ [rotenone] is the rate of change of oxygen consumption (entire flux) at low concentrations of rotenone, and ΔCI_act_ /Δ [rotenone] is the rate of change of the complex activity at low concentrations of the inhibitor. The linear rates fitted to the complex I activity and oxygen consumption data, at low concentrations of inhibitor, generated *r*^2^ values above 0.9 for the calculations of FCCs.

### Reactive Oxygen Production (ROS)

ROS was measured using dihydroethidium, a fluorescent dye which binds specifically to superoxide and hydrogen peroxide. DHE is oxidized to 2-hydroxyethidium by superoxide with an excitation wavelength of 500-530 nm and emission wavelength of 590-620 nm allowing it to be measured fluorometrically. BMECs were treated with inhibitors of complex I, III or IV. Cells were then detached using accutase and transferred to 12 mL flow cytometry tubes. Cells were washed once with PBS before resuspending in PBS + glucose (10 mM) + 20 μM DHE. Cells were incubated for 20 min at 37°C. Following incubation cells were washed twice with PBS and resuspended in PBS + glucose (10 mM). Samples were analysed using a FacsCanto flow cytometer (BD Biosciences) using the PE channel.

### Measurement of intracellular nucleotides

Nucleotides were extracted by perchloric acid precipitation. Cells were detached from 6 well plates by scraping after treatments, washed twice with medium and finally washed with PBS. Cells were homogenized in 1mL PBS in a 1 mL dounce homogenizer. Protein samples were taken before adding 1 volume of 1M perchloric acid and vortexing on ice for 5 min, every minute. Precipitated protein was spun down at 13,200rpm for 15 min at 4 degrees. Supernatant was removed (0.5 mL) and 88 μL of 2.5 M Potassium hydroxide was added for neutralization of perchloric acid. Neutralized samples were vortexed vigorously until salt formed. Samples were spun at 13,200 rpm for 15 min at 4 °C, 0.4 mL supernatant was removed and placed in a separate eppendorf tube for storage at -80 °C. Nucleotide extractions were analysed with HPLC on a C18 gemini (1.5 μm) column (Phenomenex) using an isocratic method adapted from Ingebretson *et al*. 1982. The mobile phase consisted of 220 mM potassium phosphate, 0.3 mM tetrabutylammonium hydrogen phosphate and 1% Methanol, pH 6.9. The method was run for 30min at a flow rate of 1 ml/min at 30 °C. Chromatograms were analysed used Empower pro software.

### Statistical analysis

Data and statistical analysis was performed using Graphpad prism9 (Graphpad software). One-way ANOVA was used to analyse more than two data sets with unpaired t-tests used to compare the differences between two data sets. P-value < 0.05 deemed statistically significant. *< 0.05, **< 0.01,***< 0.001, ****<0.0001.

## Results

### BMECs derived from human iPS cells exhibit high TEER values and active metabolic plasticity

To test the quality and integrity of the BBB model we measured the transendothelial electrical resistance (TEER) of differentiated human iPS cells in transwell systems (Figure 1a). TEER values of between 1500-2500 ohms/cm^2^ were routinely and consistently obtained from each differentiation (Figure 1a). Western blot analysis also confirmed the expression of the tight junction proteins: occludin, claudin-5 and ZO-1 (Figure 1a). BMECs are known to contain higher numbers of mitochondria than other endothelial cells^14^ and impaired mitochondrial function in models of stroke, aging and inflammation are associated with decreased BBB integrity^15,16–18^. Therefore we performed Seahorse XF analysis to evaluate the metabolic function of BMECs in response to specific inhibitors of mitochondrial function. A mitochondrial stress test to analyse mitochondrial metabolism in BMECs (Figure 1b) shows an oxygen consumption rate (OCR) of approximately 60-70 pmol/O_2_/min and an extracellular acidification rate (ECAR) of approximately 60 mpH/min (Figure 1b). This equates to an OCR:ECAR ratio of approximately 1:1, indicating a significant glycolytic activity. We found BMECs to be metabolically plastic, as they decrease OCR in response to the ATP synthase inhibitor oligomycin (1 μM) and increase OCR in response to the mitochondrial uncoupler FCCP (1 μM) (Figure 1b). BMECs were shown to have a substantial spare respiratory capacity (SRC) by increasing the basal rate of OCR by 30% in response to FCCP (Figure 1b). In order to balance energy deficits caused by inhibition of mitochondrial metabolism, glycolytic flux can increase to compensate for the deficit in ATP production. Here, the 14% increase in ECAR in response to oligomycin indicates that BMECs contain reserve levels of glycolytic flux to compensate for any deficits in the respiratory chain (Figure 1b).

**Figure 1:**
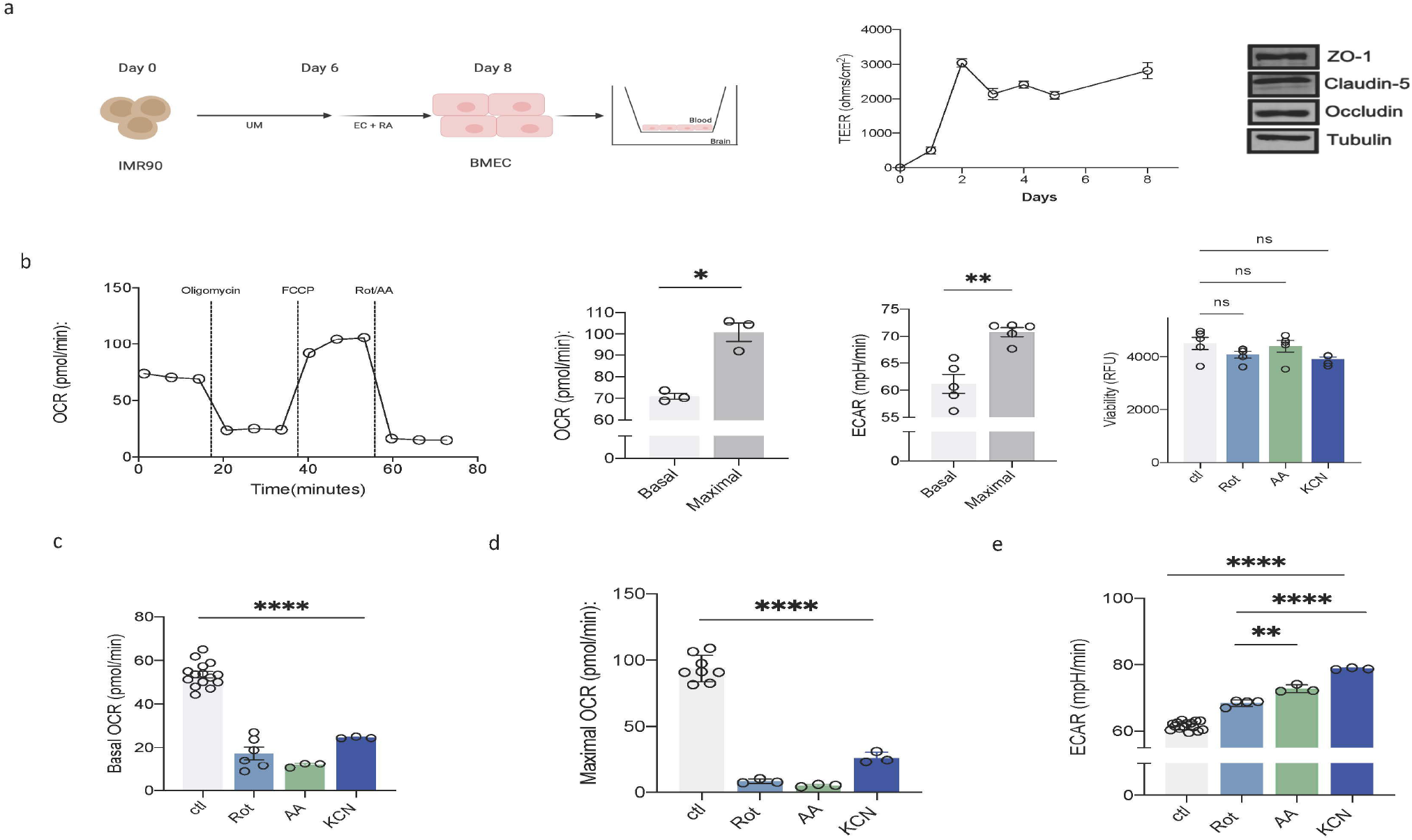
Inhibition of ETC complexes decreases oxygen consumption rates and increases glycolytic flux in BBB endothelial cells. (a) Timeline of in vitro generation of BMECs from IMR90 hIPS cells. Cells were seeded (15,000/cm) in unconditioned media (UM) and on day 6 were terminally differentiated by changing to endothelial cell (EC) culture medium plus retinoic acid. On Day 8 the cells were subcultured onto transwell filters or plates for experiments. Time course showing TEER values of BMECs sub-cultured onto transwell filters. Expression of tight junction proteins occludin, ZO-1 and claudin-5 in fully differentiated BMECs (b) Oxygen consumption rate (OCR) trace from Seahorse XF analysis of fully differentiated BMECs. FCCP uncoupling allows maximal OCR to be measured. Maximal ECAR is indicative of increases in glycolytic flux. (c) Basal OCR in response to ETC inhibitors, Rotenone (1 μM, Rot), Antimycin A (2 μM, AA), Potassium Cyanide (1.5mM, KCN) (d) Maximal OCR in response to same ETC inhibitors (e) Glycolysis rates in response to same ETC inhibitors. All results are from 3 independent experiments. One way ANOVA and unpaired t-tests were used to calculate statistical significance with a P-value < 0.05 deemed statistically significant. *< 0.05,**< 0.01,,****<0.0001.

The control of metabolism by ETC complexes in BMECs was studied using specific inhibitors of CI (rotenone, 1 μM), CIII (antimycin A, 2 μM) and CIV (KCN, 1.5 mM), at concentrations that did not affect cell viability (Figure 1b). Effects of ETC complex inhibition on OCR and ECAR are shown in Figures 1c-e. Inhibition of CI, CIII or CIV significantly decreased OCR in BMECs over an 18-minute time period (Figure 1c). Maximal OCR and the SRC of BMECs were also significantly compromised in response to CI, III or IV inhibition (Figure 1d). Interestingly, there was a corresponding increase in glycolysis (Figure 1e), most likely to maintain ATP levels and mitochondrial membrane potential through the reversal of ATP synthase and hydrolysis of glycolytic ATP^19^.

### BMECs exhibit a highly fused and tubular mitochondrial network with mitochondrial membrane potential and dynamics sensitive to ETC inhibition

In response to energetic stress and a need to maximize ATP production mitochondria fuse in order to share matrix components such as subunits of ETC complexes and mtDNA^20^. This allows for the maintenance of the ETC and prevention of mitochondrial dysfunction, pathological ROS, dissipation of ΔΨ_m_, and apoptosis^21^. Here, using confocal microscopy we show that BMECs exhibit a highly fused and reticular mitochondrial network (Figure 2a), indicative of enhanced mitochondrial fusion. To ascertain the effects of ETC inhibition on ΔΨ_m_, cells were stained with TMRM and real time live cell imaging of TMRM stained BMECs was performed over <u>60</u> min following treatment with Rot (1 μM), AA (2 μM) or KCN (1.5 mM) (Figure 2b). Inhibition of CI, CIII or CIV significantly reduced ΔΨ_m_ (Figure 2c). Next, we examined the effect of mitochondrial inhibition on proteins involved in mitochondrial fission-fusion dynamics. The long form of the inner membrane fusion protein OPA-1 (L:OPA1) was reduced in response to CI-V inhibitors and this correlated with an increase in the short form, S:OPA1. Complete loss of L:OPA1 in response to the mitochondrial uncoupler FCCP was also observed (Figure 2d). Mitochondrial fission factor protein (Mff) acts as a docking site for Drp1, the main driver of mitochondrial fission, and the expression of this Mff was increased in response to CI-V inhibition (Figure 2d). These results show that inhibition of CI, CIII and CIV significantly reduce mitochondrial function, leading to a substantial reduction in ΔΨ_m_ and increased mitochondrial fission.

**Figure 2:**
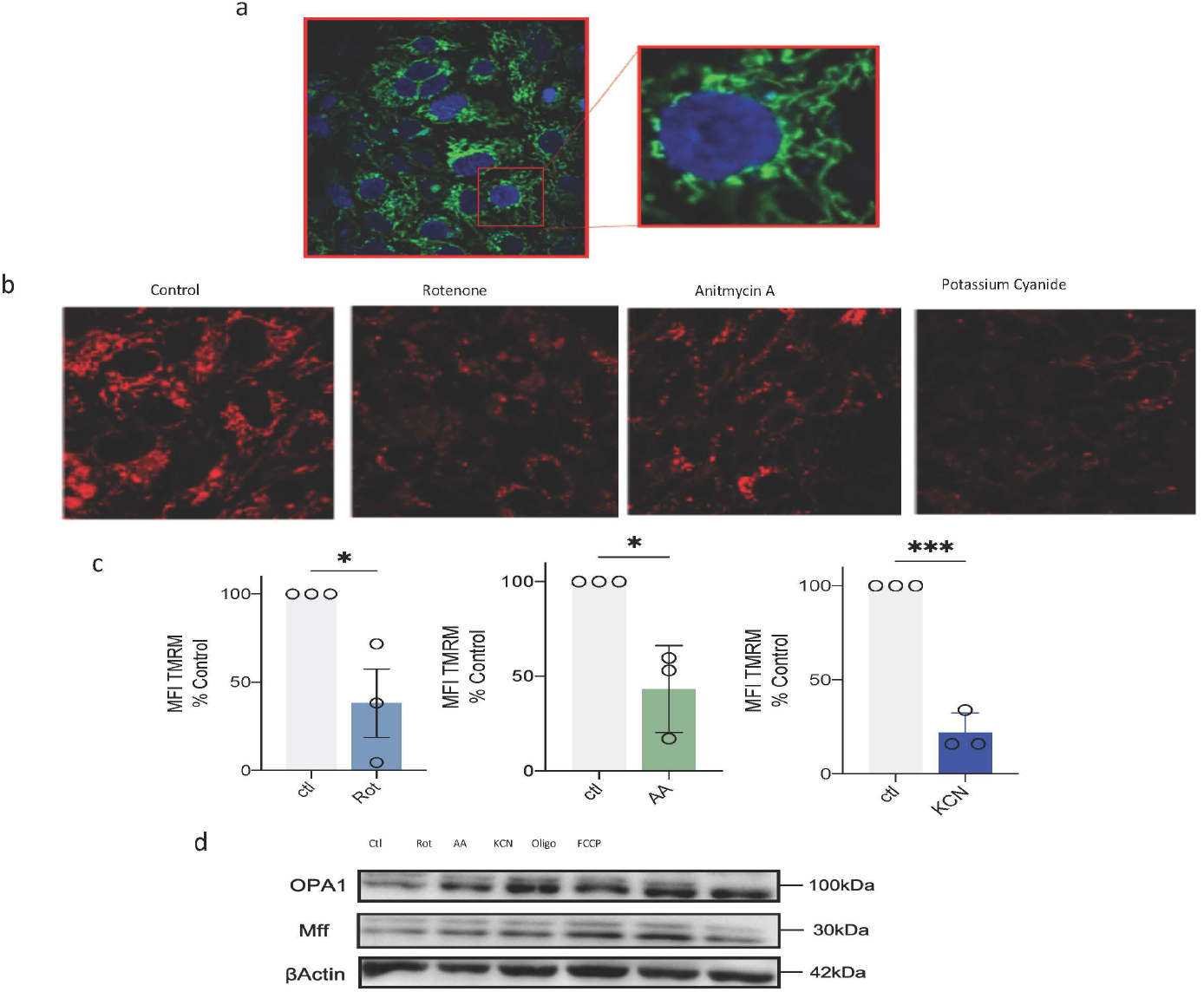
BMECs exhibit a highly fused and tubular mitochondrial network with mitochondrial membrane potential and dynamics sensitive to ETC inhibition. (a) Confocal image of mitochondrial network in BMECs, with mitochondria labelled green and nuclei labelled blue (b) Confocal images of mitochondria stained with TMRM treated with Rotenone (1 μM, Rot), Antimycin A (2 μM, AA), Potassium Cyanide (1.5 mM, KCN) (c) Mean fluorescence intensity (MFI) of mitochondria stained with TMRM treated with same ETC inhibitors for 60 min represented as percentage of control (d) Western blot analysis of OPA1 and Mff expression in BMECs after treatment with ETC inhibitors for 60 min. (a), (b) and (c) are representative of 3 independent experiments represented as Mean±SEM as a percentage of control MFI. Unpaired t-tests were carried out to deem statistical significance with a P-value < 0.05 considered statistically significant. *< 0.05,***< 0.001.

### Complex I has a high degree of control over BBB permeability and oxidative phosphorylation in BMECs

As mitochondrial CI dysfunction is a hallmark of aging and neurodegenerative disorders such as PD and AD^22^, we investigated the effect of inhibition of ETC complex activities on the integrity of the BBB in a transwell system. TEER values rapidly and significantly decreased by 60% when CI activity was inhibited with 1 μM rotenone. However, treatment with high concentrations of AA (2μM) or KCN (1.5mM) did not decrease TEER values over this time period (Figure 3b) even though OCRs were significantly reduced similar to rotenone-inhibited cells (Figure 1c-d). To rule out the possibility that rotenone inhibits microtubule formation in BMECs, we used an alternative complex I inhibitor, piericidin A, not known to effect microtubule dynamics. Following introduction of 5μM piericidin A, TEER values were immediately reduced by approximately 40% (Figure 3a). Direct addition of an inhibitor of microtubule assembly, nocodazole, resulted in no significant changes in TEER values (Figure 3b). These results indicate that inhibition of CI activity has a critical role in controlling BBB permeability potentially through maintenance of mitochondrial and bioenergetic homeostasis.

**Figure 3:**
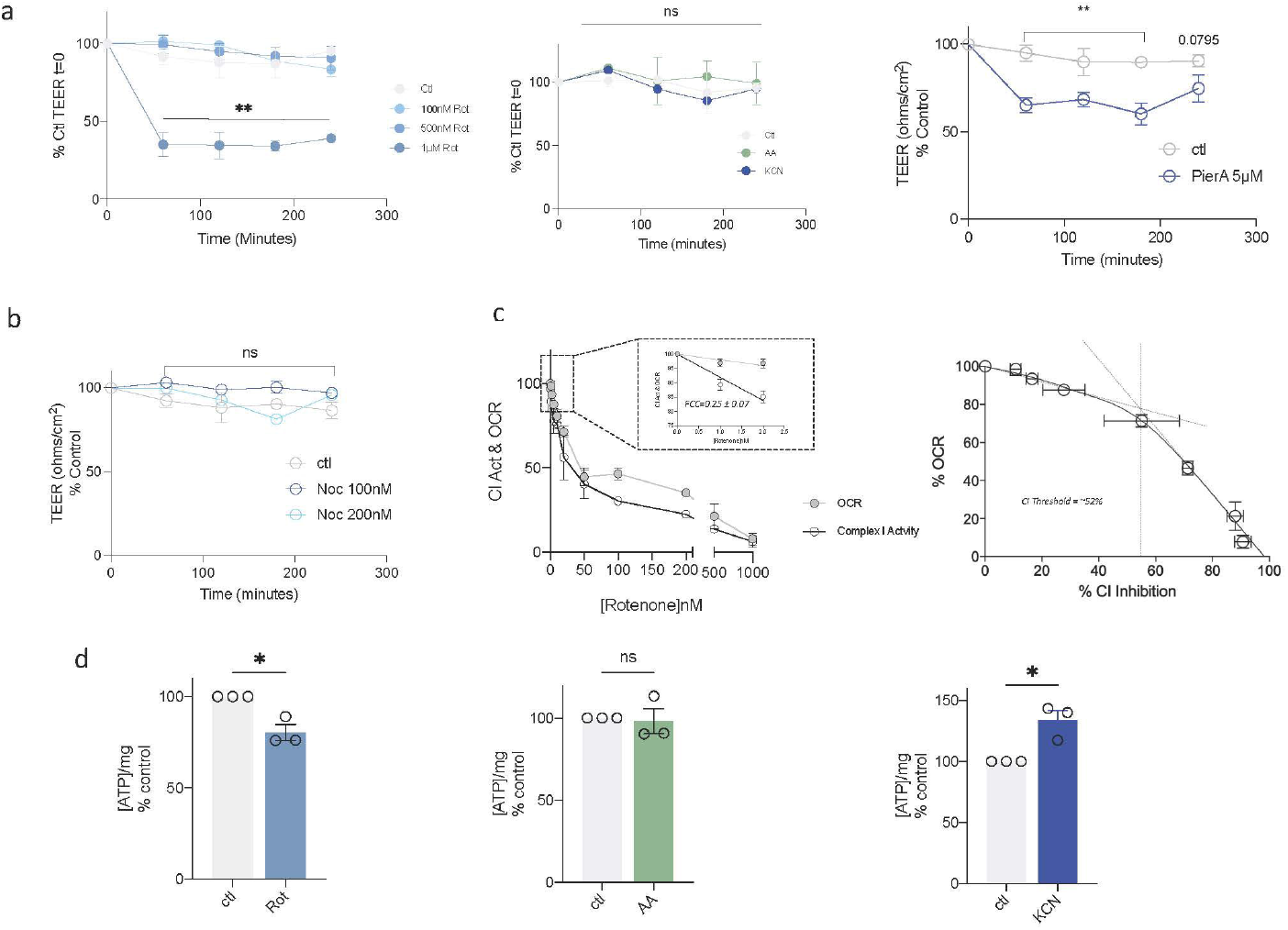
Overcoming complex I energy thresholds in BMECs decreases TEER and ATP. (a) TEER time course after treatment with increasing concentrations of Rotenone (100nM-1μM Rot), Antimycin A (2μM, AA) and Potassium Cyanide (1.5mM, KCN), Piericidin A (5μM) (b) TEER time course after treatment with nocodazole (100nM-500nM) (c) Oxygen consumption rates measured by Seahorse XF analysis and Complex I activity measured spectrophotometrically by the oxidation of NADH at 340nm. Also shown is Complex I inhibition threshold curve represented as the decrease in OCR as a function of increasing Complex I inhibition (d) ATP levels after treatment in ETC inhibitors for 1 h presented Unpaired t-test and paired t-test was carried out to deem statistical significance with a P-value < 0.05 considered statistically significant. *< 0.05,**< 0.01.

CI is a primary entry point for electrons into the ETC and its dysfunction is associated with disorders such as PD and AD where BBB integrity is compromised. We therefore examined the level of control CI has over oxidative phosphorylation in BMECs. We used Metabolic Control Analysis (MCA) to quantify the level of control CI has over BMEC respiration. By measuring CI specific activities and OCRs during a rotenone titration, we calculated a flux control coefficient of 0.25 ± 0.07 for CI in BMECs (Figure 3c). This flux control coefficient indicates that slight perturbations in CI activity immediately control 25% of mitochondrial respiration in BMECs. We further used this data to calculate an inhibition threshold of 55% for CI in BMECs, indicating the level of CI inhibition BBB cells can withstand before large changes in respiration are observed (Figure 3c). HPLC analysis of BBB cells showed that CI inhibition significantly reduced ATP levels after 1h whereas no effects were seen with CIII inhibition. Inhibition of CIV activity significantly increased ATP levels (Figure 3d), likely a consequence of increased glycolytic flux (Figure 1e).

On interpreting the results of the MCA analysis it is evident that while CI is a critical control point for immediate control of metabolic flux in BBB cells, it can withstand large levels of inhibition before oxidative phosphorylation in these cells is severely compromised.

### Restoration of BBB integrity follows transient increase in complex I-induced permeability

To further investigate BBB permeability over an extended period of time we synthesized fluorescein isothiocyanate (FITC) labelled bovine serum albumin (BSA) protein (BSA-FITC) and examined its transport across the endothelial monolayer during 16 h of ETC inhibition. The transport of BSA-FITC across the BMEC monolayer from the apical to the basolateral chamber was significantly increased only in response to CI inhibition (Figure 4a) and the initial CI related loss in TEER was fully recovered after 16 h (Figure 4b). This indicated that CI inhibition beyond the 55% inhibition threshold caused immediate disruption to the BBB permeability, facilitating entry of BSA-FITC across the BMEC monolayer. While inhibition of CIII or IV had no immediate effect on transport of BSA-FITC (Figure 4a) or BBB permeability, TEER values increased over the extended time period (Figure 4b). Even though respiration was inhibited during the 16 h period (Figure 5) ATP levels were fully restored during incubation with all three ETC inhibitors (Figure 4c), however, there were no effects on ADP, AMP or NAD levels (Figure 6).

**Figure 4:**
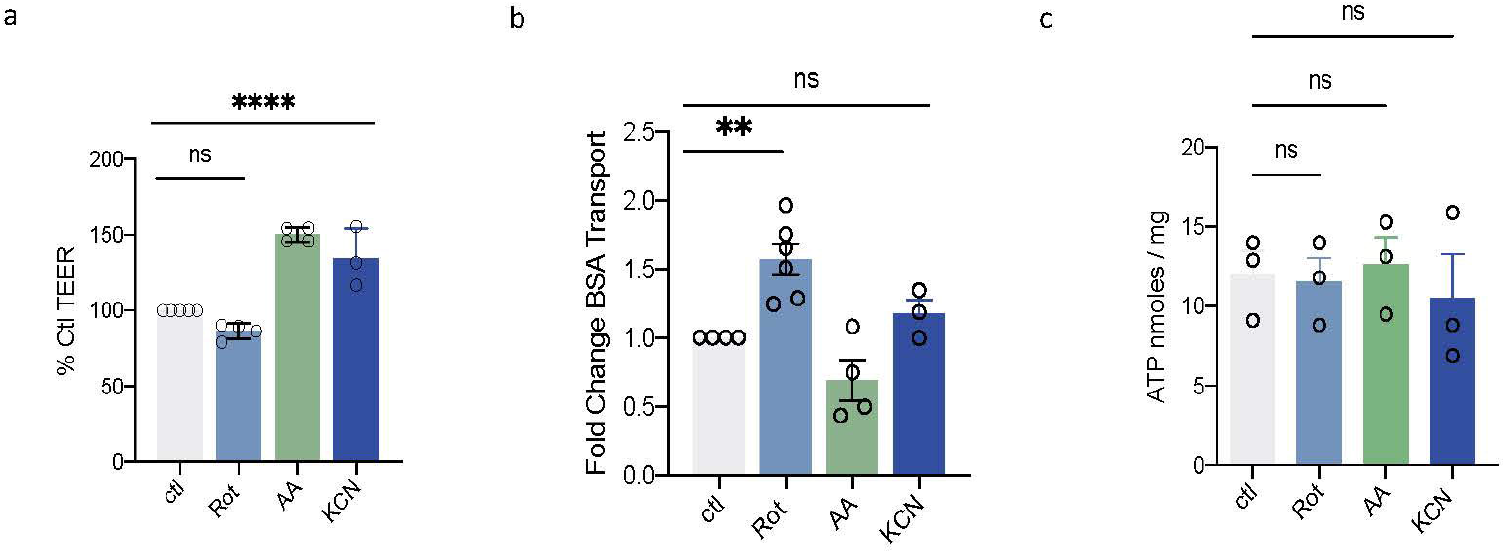
Inhibition of Complex I causes increased blood brain barrier permeability and transport across BMEC monolayers. (a) TEER value after 16 h treatment with ETC inhibitors represented as percentage of control TEER value (b) Transport of BSA-FITC across BMEC monolayers over 16 h after treatment with Rotenone (Rot, 1 μM), Antimycin A (AA, 2 μM) and Potassium Cyanide (KCN, 1.5mM). Data shown from 4 independent experiments (c) Concentration of ATP per mg of protein after 16 h treatment with ETC inhibitors represented as a percentage of the control concentration. One way ANOVA and unpaired t-tests were used to calculate statistical significance with a P-value < 0.05 deemed statistically significant. *< 0.05,**< 0.01,****<0.0001

**Figure 5:**
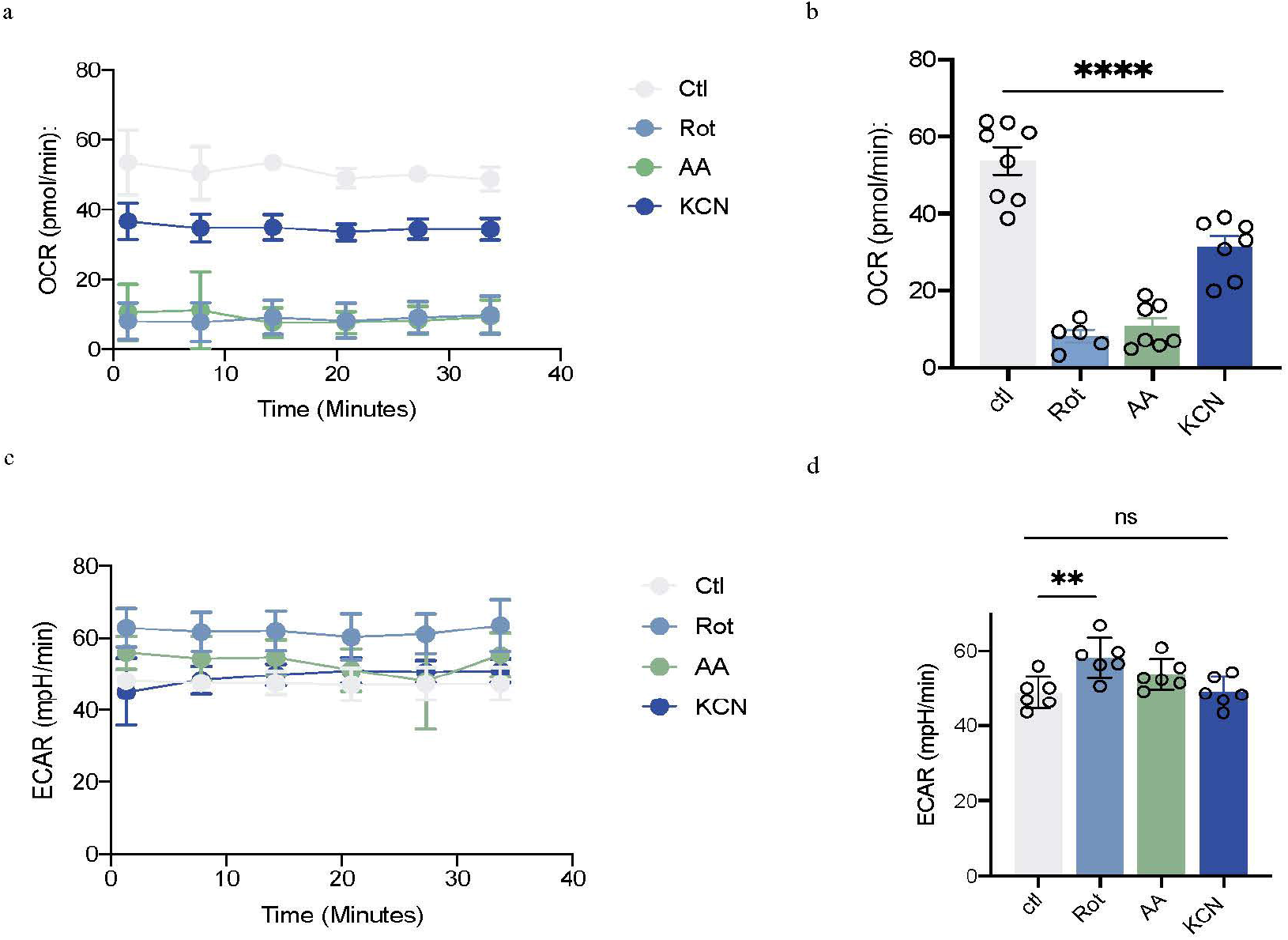
SeaHorse analysis of respiration and glycolytic flux following extended incubation with ETC inhibitors. BMECs treated with ETC inhibitors for 16 h (Rot-1μM, AA-2μM, KCN-1.5mM). Control cells treated with vehicle (0.1% EtoH). OCR (a&b) and ECAR (c&d) were analysed using a 96 well seahorse XF flux analyzer. One way ANOVA were used to calculate statistical significance with a P-value < 0.05 deemed statistically significant.**< 0.01,****<0.0001.

**Figure 6:**
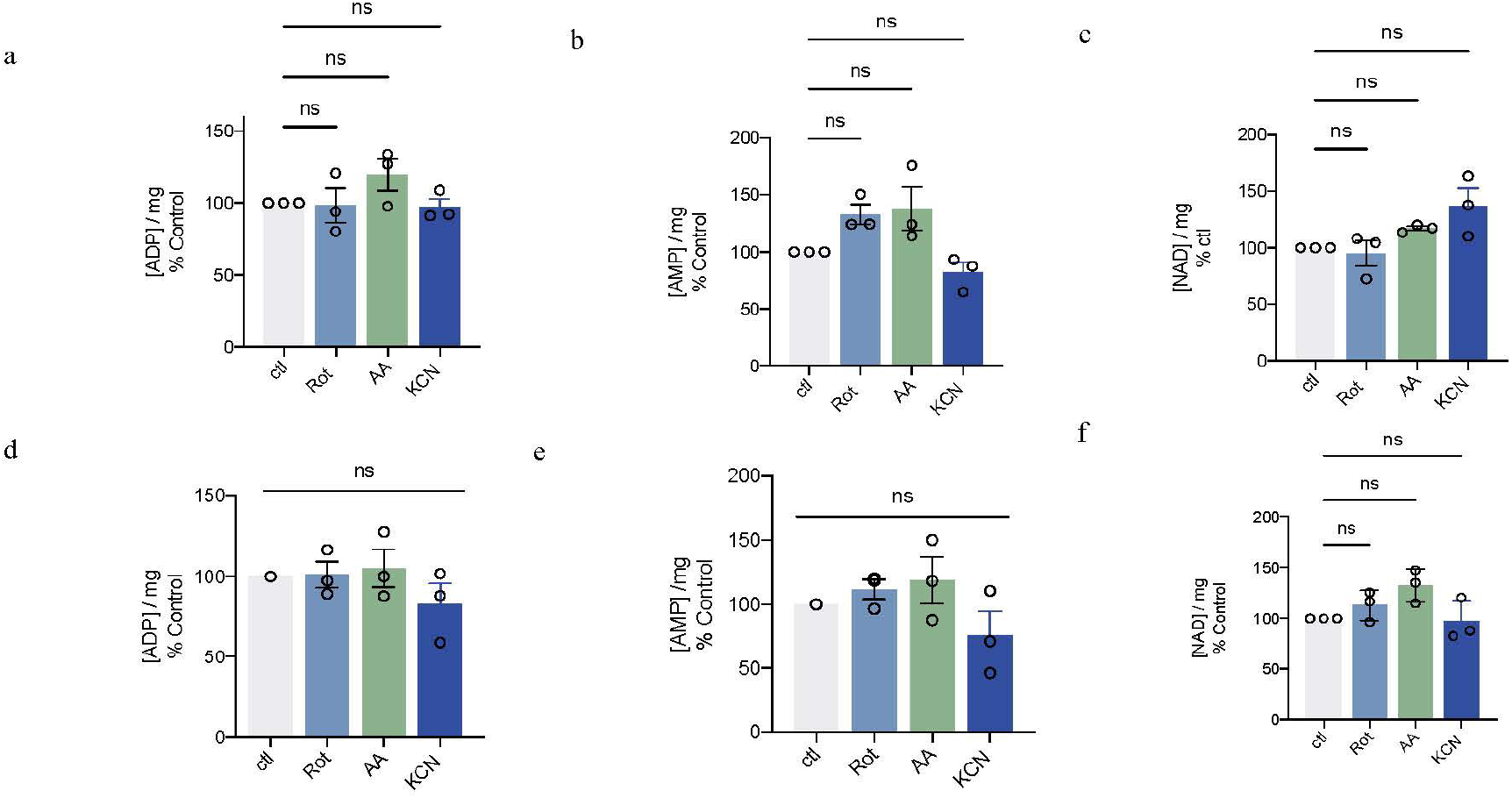
Inhibition of ETC complexes does not change the level of ADP, AMP and NAD in BMECs. (a-c) ADP, AMP, NAD levels after 1 h treatment with ETC inhibitors (Rot-1 μM, AA-2 μM, KCN-1.5 mM) represented as a percentage of control (d-f) ADP, AMP, NAD levels after 16 h treatment with ETC inhibitors represented as a percentage of control. All data representative of 3 independent experiments. One way ANOVA were used to calculate statistical significance with a P-value < 0.05 deemed statistically significant.

We then tested the effect of ETC inhibition on ROS production in BMECs using the superoxide probe dihydroethidium (DHE) and found no differences in ROS produced at CI or CIII in response to respective ETC inhibitors over 4 h that could account for the differential BBB permeability effects (Figures 7a-b). Similarly, following a 16 h time-period no differences in ROS formation at CI or CIII were observed (Figure 7c-d) that could account for differential BBB permeability effects.

**Figure 7:**
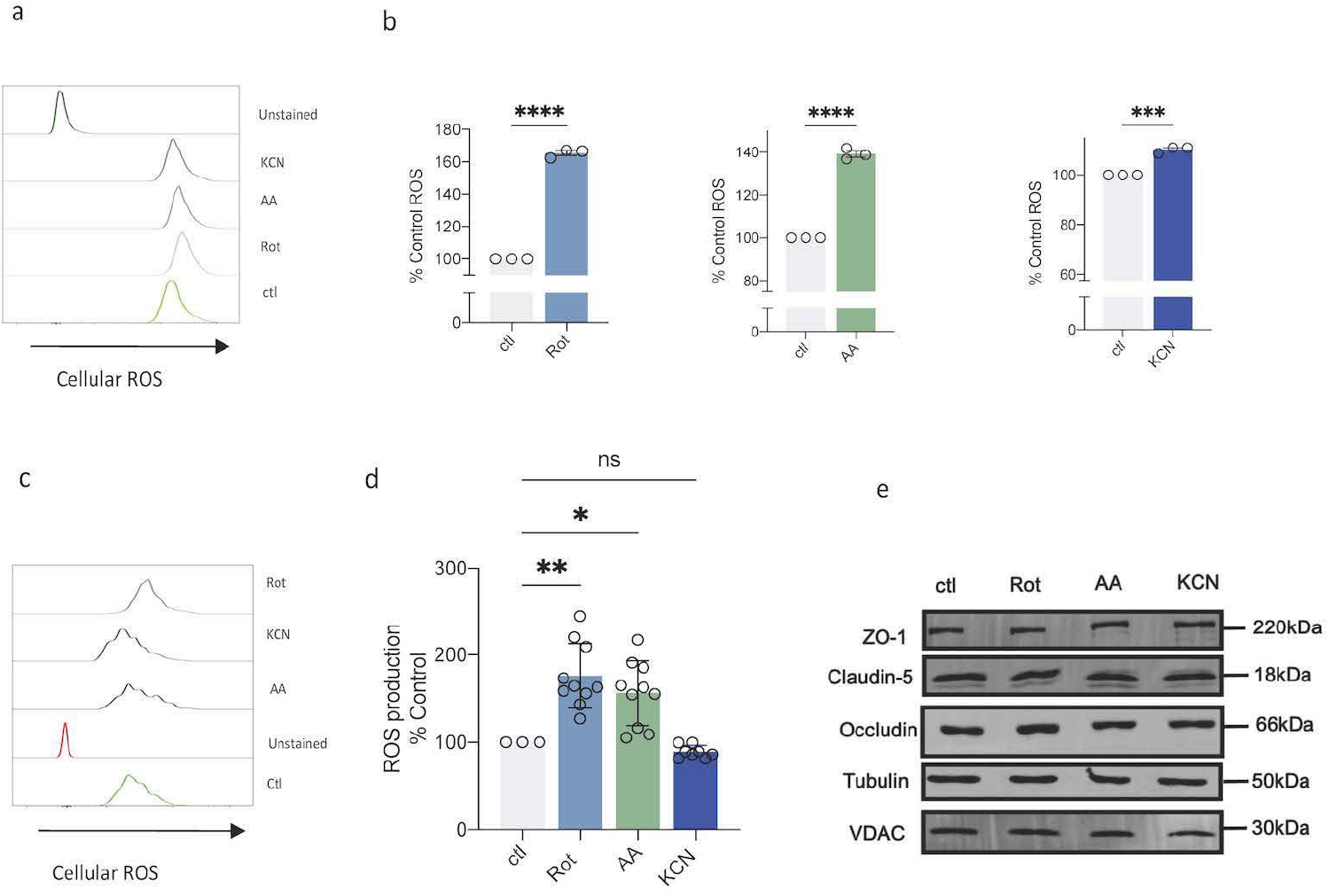
ETC inhibition increases ROS production in BMEC monolayers. (a) Flow cytometry histograms of DHE mfi representing ROS production in BMECs treated with ETC inhibitors for 1 h (b) ROS production measured by flow cytometry using DHE represented as percentage of control in response to different ETC inhibitors for 1 h (c) Flow cytometry histograms of DHE MFI measuring ROS production over 16 h (d) Cellular ROS values 16 h post treatment with ETC inhibitors represented as percentage of control (e) Western blot analysis of tight junction proteins after 16 hour treatment with ETC inhibitors. Data shown from 4 independent experiments except (e) which is representative. All data represented as the Mean ± SEM. One way ANOVA and unpaired t-tests were used to calculate statistical significance with a P-value < 0.05 deemed statistically significant. *< 0.05,**< 0.01,***< 0.001, ****<0.0001.

We also performed western blot analysis of tight junction proteins in BMECs treated with inhibitors of CI, III and IV over the 16 h time course. We observed no change in the expression of the tight junction proteins occludin and claudin-5 in BMECs treated with rotenone, antimycin A or KCN (Figure 4f) that could account for the CI-related control of BBB permeability. However, increased expression in ZO-1 in response to CIII and IV inhibition was observed (Figure 4f), correlating with the observed increase in TEER value and increased glycolytic flux under these conditions.

## Discussion

In this study we used a Metabolic Control Analysis approach to highlight the high level of control which mitochondrial complex I has over respiration in human induced pluripotent stem cell derived BMECs. We identify a role for mitochondrial complex I in controlling permeability and transport of protein across an *in vitro* BBB system.

BMECs are specialized endothelial cells that form the cerebral vasculature and generate an effective barrier between the blood and brain. Maintenance of BBB homeostasis requires energy in the form of ATP. Mitochondria contribute to the production of ATP, as well as several other functions including, calcium homeostasis, ROS production and the induction of programmed cell death ^23,24^. BMECs obtain most of their ATP from glycolysis, however there is limited knowledge on the contribution of mitochondria to the maintenance of bioenergetic status in these cells.

BMECs are known to contain higher levels of mitochondria than any other endothelial cells in the body^14^. Recent studies have highlighted that impaired mitochondrial function in models of stroke and inflammation cause increased BBB permeability, and deletion of mitochondrial specific proteins causes loss of BBB integrity ^15,16,25^. We examined how specific complexes of the ETC control the global flux of mitochondrial oxygen consumption and ATP synthesis in BMECs. As expected BMECs derived from human iPS cells exhibited active mitochondrial respiration which was sensitive to ETC inhibitors. The higher mitochondrial content in BMECs was reflected in the substantial (30%) SRC. Complex I, III and IV inhibition all decreased basal and maximal respiration in BMECs indicating the dispersion of control over the flux of respiration. This increase in glycolytic flux could aid in maintaining energy homeostasis when mitochondria are impaired in the BBB, with BMECs holding a glycolytic reserve ready to respond to energetic stress such as ETC inhibition, decrease in energy sources or infection. Depending on the site in the ETC that was inhibited, there was a corresponding increase in glycolysis. Inhibition of CI activity caused the least significant increase, with CIII and CIV inhibition causing a more rapid glycolytic response, as indicated by increased ECAR. This indicates that the glycolytic flux in BMECs is sensitive to ETC inhibition and depending on the site of inhibition on the ETC, glycolysis can respond differentially. This cross-talk between glycolytic flux and mitochondrial respiration may be important for maintaining energy homeostasis in BMECs and preventing BBB dysfunction. Similarly, in another BBB cell culture model involving hCMEC/D3 cells, a reduction in OCR values following loss of cardiolipin was counter-balanced by increased glycolytic rates ^26^.

Redox homeostasis is dependent on ΔΨ_m_ and is critical for the maintenance of BBB integrity, highlighting the need for a well organised mitochondrial network. The results in Fig 2 support previous findings that mitochondrial dynamics have been shown to effect BBB integrity and permeability. Recently, LPS induced BBB dysfunction associated with sepsis activated Drp-1, the main executioner of mitochondrial fission, leading to a decrease in ΔΨ_m_, an increase in BBB permeability and a loss of tight junctions has been reported ^25^.

Mitochondrial energy thresholds have been characterized for CI in several tissues^27^ and such thresholds may underlie neuronal survival and function in the brain^28^. CI has a low inhibition threshold of 12-25 % in mitochondria located in the nerve terminal^29^, indicating that a small loss of CI activity in these mitochondria will rapidly lead to compromised ATP generation and excessive release of excitotoxic glutamate^30^. However, non-synaptic mitochondria can withstand much higher levels of inhibition (72%) before respiration and ATP synthesis are compromised^31^. Our study shows that BMECs possess high CI inhibition and energy thresholds and when these are overcome a reduction in TEER values will present. A role for reduced mitochondrial respiration in compromising the integrity of the BBB through the actions of MiR-34a, a microRNA shown to have several mitochondrial targets, has been identified^16^. MiR-34a was shown to dose dependently inhibit respiration with corresponding decrease in ATP production and cytochrome c levels, increasing blood brain barrier permeability^16^.

These data indicate the involvement of CI dysfunction in controlling BBB permeability, potentially linking it to disease with observed BBB dysfunction, where CI dysfunction is also present. A recent study showed increased BBB permeability and altered tight junctions present following deletion of Crif1, an essential protein in assembly of the ETC^32^. Specific downregulation of the CI subunit NDUFA9, a core CI subunit involved in the oxidation of NADH and commonly associated with mitochondrial disorders, which is downregulated in the AD brain^32^. This indicates that CI could contribute to BBB dysfunction in neurological disorders such as PD and AD where a reduced CI activity is present.

Similar differential responses to inhibition of ETC activities have been identified in tumorigenesis and angiogenesis where complex III was found to control hypoxic activation^33,34^, whereas complex I regulates organismal survival and motor control in a mouse model of mitochondrial dysfunction^35^. Our results suggest that downstream ETC inhibition at complexes III or IV in BMECs induces a more significant glycolytic response that maintains ATP levels. This response is not present when complex I is inhibited, corresponding to decreased ATP levels and increased BBB permeability. However, these effects are transient and the BMECs recover bioenergetic homeostasis and permeability resistance over an extended time period. Mitochondrial supercomplexes involving differential stoichiometries of ETC complexes that form respirasomes, have been identified across a range of cells ^36^. The NADH-respirasome and CoQ ^37^ have yet to be fully characterised in mitochondria in BBB endothelial cells, and the organisation of ETC complexes may play a role in the differential regulation of BBB permeability, as observed in this study.

We have shown that under our experimental conditions complex I inhibition causes decreased mitochondrial respiration, decreased ΔΨ_m_ and altered mitochondrial dynamics in BMECs. Inhibition of complex I also decreased TEER values in *in vitro* BBB systems, with a corresponding increase in transport of protein across BMEC monolayers. These data contribute to the current understanding of the role of mitochondria in the cerebral vasculature. The BBB is a complex system, and its dysfunction can exacerbate cerebral dysfunction leading to enhanced neuronal destruction in diseases such as PD, AD and stroke.

Safeguarding complex I related mitochondrial function may offer a novel therapeutic strategy for the maintenance of BBB integrity. The selective complex I control of BMEC permeability may also provide a target for enhancing transport of therapeutics and nanoparticles across the BBB^38–40^.

## Acknowledgements

Funding is gratefully acknowledged from the EU Marie Curie Initial Training Network (FP7 People: Marie-Curie Actions), Project No. 608381.

## Author Contributions

GMD performed all the experiments and co-wrote the manuscript. EJ synthesised the BSA-FITC. JMH co-designed the experiments. GPD designed, oversaw the study, and co-wrote the manuscript. All authors read and approved the final manuscript.

